# Valence-dependent synaptic plasticity drives approach and avoidance behavior in social context

**DOI:** 10.1101/2022.11.08.515650

**Authors:** Pedro Espinosa, Benoit Girard, Mattia Lucchini, Federica Campanelli, Valentina Tiriticco, Lucy Mohrhauer, Camilla Bellone

**Affiliations:** University of Geneva, Department of Fundamental Neuroscience; CMU, 1 rue Michel-Servet, 1211, Geneva, Switzerland

## Abstract

The nature of social interactions determines engagement or avoidance with conspecifics. Here, we explore the circuit mechanisms that guide approach or avoidance behavior in mice based on the valence of previous social interactions. We identify a novel circuit connecting D1 receptor expressing neurons of the anterior insular cortex (AIC) to D1R expressing neurons of the nucleus accumbens (NAc). These cells become active during social interactions in a valence-dependent manner. Lower frequency patterns encoded appetitive interactions, while aversive interactions led to higher activation. These activity patterns elicited distinct forms of synaptic plasticity in the accumbal target neurons, which were causal for subsequent approach or avoidance behavior. Our results unravel the synaptic mechanisms instructing behavior after the social interaction of opposite valence.

## Main Text

Understanding how individuals navigate social interactions is crucial for the survival of all species, relying on the ability to discern between positive and negative experiences, a concept known as social valence (*1*). Following an initial interaction, information gained helps an individual decide whether to approach or avoid a particular individual in the future. This process has been extensively studied, nevertheless the brain circuits contributing to assigning valence and learning from the experience to take the appropriate decision remain elusive. Within the mesolimbic system, the Nucleus Accumbens (NAc) contributes to the learning process by updating future behavior by guiding decisions (*2*). Although the NAc has been implicated in both rewarding and aversive social experiences (*3*-6) how it discriminates between social valence is still a matter of debate. Here we design a paradigm that allows us to investigate the neuronal mechanisms that occur when an individual enters in contact with another to assign the valence and learn to approach or avoidance behavior based on the experience.

### Juvenile and CD1 interactions are classified as rewarding or aversive experiences

We first exposed male mice to a sex-matched juvenile or an adult aggressive CD1 mouse for 15 minutes, using an inanimate object as the control (Fig. 1A). The following day, we placed the same stimuli in an enclosure within a corridor arena. We measured the time the experimental mouse spent near that enclosure during the pre-test and post-test phases. Mice showed increased interaction time with juvenile mice (indicating approach behavior) and decreased time spent near the aggressive CD1 mice (indicating avoidance behavior) during the test phase (Fig. 1, B and C). To confirm that avoidance was induced by a negative experience, we exposed mice to a non-aggressive CD1 mouse. In this case, no avoidance behavior was observed (fig. s1, A to C). Since aggression is necessary to trigger avoidance, we hypothesized that contact bouts during the specific interaction with a conspecific are responsible for inducing approach and avoidance behaviors. Therefore, we manually classified contact interaction bouts between the mice on day 1 as either positive or negative (Fig. 1, D and E). Interactions with juvenile mice were predominantly characterized by positive bouts, whereas most bouts with CD1 mice were negative (Fig. 1, F and G). To extend our analysis across the entire dataset, we employed Simple Behavioural Analysis (SimBA) (*7*). Based on this comprehensive data analysis, we classify the experience with the juvenile mouse as rewarding and the experience with the CD1 mouse as aversive (Fig. 1H).

**Fig. 1.**
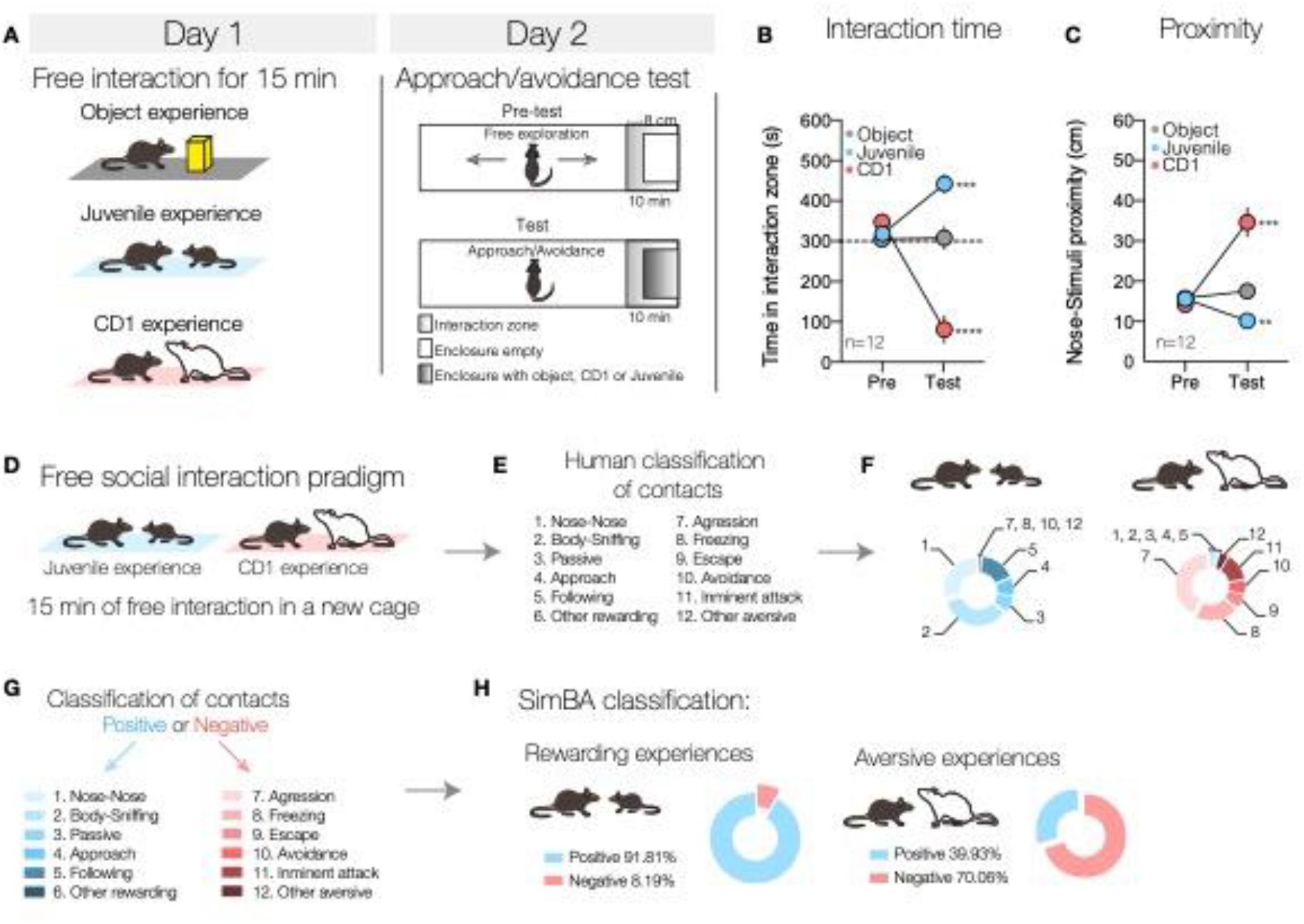
Juvenile and CD1 experiences induce learned approach and avoidance behavior. (**A**) Mice were exposed to an object, a juvenile, or CD1 mice for a 15-minute period of free social interaction. After 24 hours, the animals were placed in a new arena where, following 10 minutes of free exploration with an empty enclosure, an enclosure containing the stimuli from the previous day was introduced. (**B**) Time spent in the interaction zone on day 2, with *** indicating p = 0.0002, **** denoting p < 0.0001, and p = 0.9130 for interactions with the object. (**C**) Proximity to the stimulus mouse on day 2, measured as the distance between the mouse’s nose point and the center of the enclosure; ** indicates p = 0.0013, *** represents p = 0.0001, and p = 0.3020 for the object. (**D-E**) Schematics showing the manual labels assigned to contacts and their subsequent quantification (F). (**G**) Classification of contacts into positive and negative categories in preparation for training SimBA. (**H**) Classification of contacts by SimBA.

### D1-expressing neurons of Lateral NAc shell (LNAc) encoded proximity regardless of the valence

To study how D1R-containing MSNs compute social valence, we focused on LNAc. It has been shown that subdivisions of the NAc can exert opposing effects on motivated behavior, with the lateral portion of the NAc shell encoding rewarding and aversive experiences (*6*). We used miniature microscope to record the calcium (Ca^2+^) activity of individual D1-MSNs during rewarding and aversive social experiences (Fig. 2A). Ca^2+^ transients increased during social experiences (Fig. 2, B and C). Furthermore, we found that the Ca^2+^ activity increased when social stimuli were in proximity to the experimental mouse (Fig. 2, D to F and fig. s2, A to D), regardless of the identity of the stimulus and of the contacts (Fig. 2, E and F, fig s3). Notably, 74.6% of neurons exhibited increased activity during rewarding and aversive experiences (Fig. 2G). Since there were no discernible differences in z-score activity between the two experiences, we analyzed of the frequency domain using Fast Fourier Transform (FFT) to detect any variations in frequencies (Fig. 2H). However, no distinctions were observed between rewarding and aversive social experiences (Fig. 2, I to K). Thus far, our findings indicate that LNAc D1-MSNs encode conspecific proximity regardless of the valence.

**Fig. 2.**
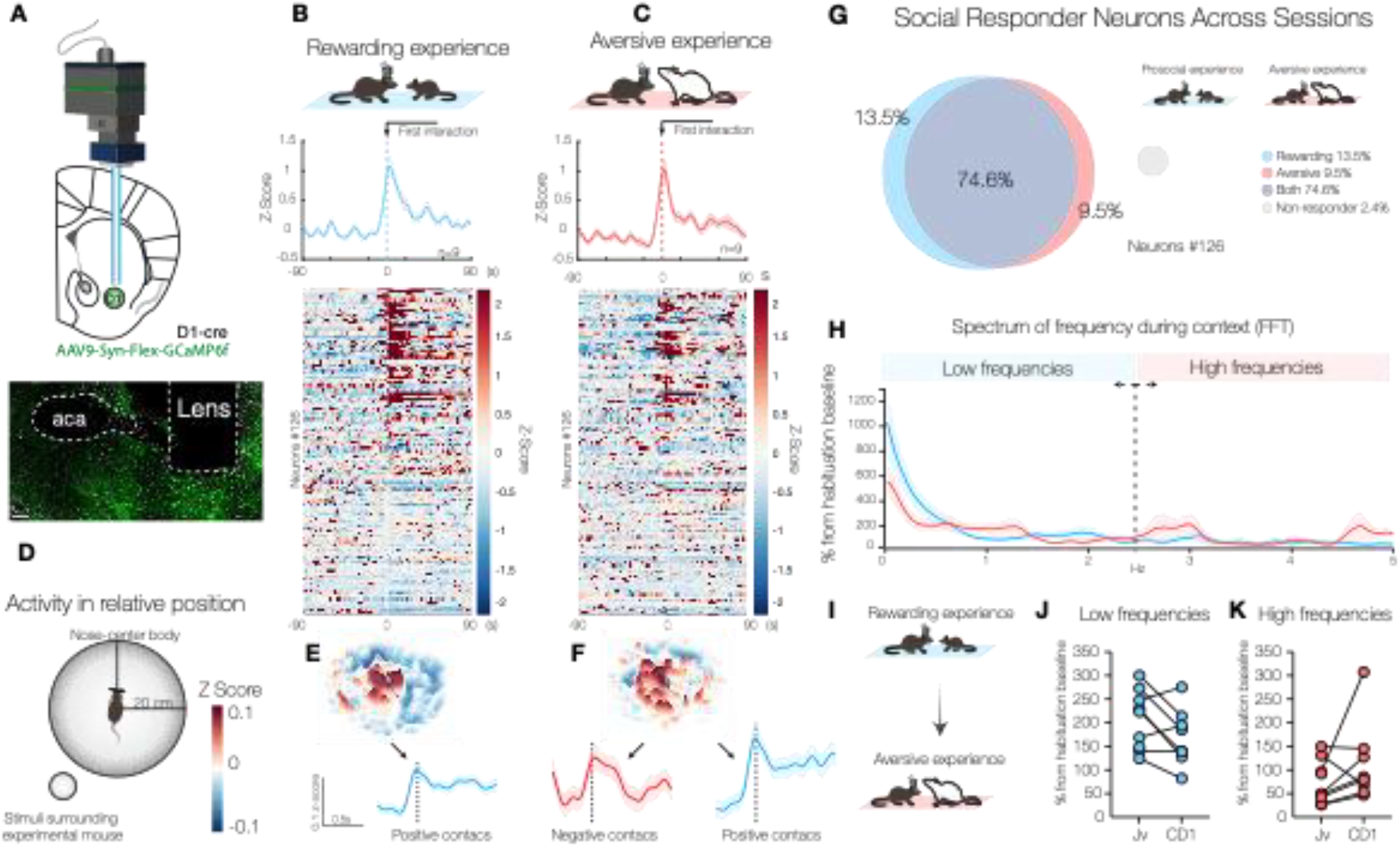
D1-MSNs encode proximity to social stimuli, independent of experience valence. (**A**) Schematic of injection and implantation sites (Scale 250uM). (**B**) Average activity during a rewarding experience aligned with the first interaction. The heatmap represents the activity of individual neurons (each row represents one neuron). (**C**) Average activity during an aversive experience aligned with the first interaction. The heatmap represents the activity of individual neurons (each row represents one neuron). (**D**) Schematic of a 3D heatmap representing the calcium activity in position relative to the stimuli. (**E**) Heatmap showing the activity in positions relative to juvenile stimuli. (**F**) Heatmap showing the activity in positions relative to CD1 stimuli. (**G**) Proportion of neurons that respond to social stimuli. (**H**) Spectrum of frequencies derived from a Fast Fourier Transform (FFT). (**I**) Schematic showing the animals’ first and second exposures. (**J**) Quantification of the low-frequency spectrum between the two exposures (0-2.5Hz). (**K**) Quantification of the high-frequency spectrum between the two exposures (2.5-5Hz). All analysis were performed on the same set of neurons that were tracked across sessions for all panels.

### AIC-NAc projecting neurons encoded the social context

We used a neural activity-dependent labeling approach to investigate the inputs that drive D1-MSN activity during rewarding and aversive social experiences. We injected Fos-cre-ERT2 mice in the LNAc with a retro-AAV-FLEX-tdTomato virus and, four weeks later, exposed the mice to non-social, rewarding, or aversive stimuli for 15 minutes, followed by tamoxifen administration (fig. s4A). We identified neurons activated in different rostro-caudal subdivisions of the insular cortex (anterior insular cortex - AIC, medial insular cortex - MIC, and posterior insular cortex - PIC) (*8*) and in the amygdala (fig. s4, B to C). The insular cortex has an integrative function, plays an essential role in salience processing and attention, and gates executive control (*9*). These functions are implicated in social behavior and valence processing (*10, 11*). By injecting a retrograde rabies virus into the LNAc, we found that only the anterior section of the insula projects monosynaptically to the D1-MSNs in the lateral part (fig. s5, A to D). To identify the identity of socially active neurons in the AIC that project to the LNAc, we used RNA scope in situ hybridization to double label c-Fos and Drd1 mRNA. We observed that 88.16% of juvenile-activated neurons and 81.4% of CD1-activated neurons were Drd1-positive (fig. s6, A and B). We confirmed these results using a bicolor virus approach, injecting AAVrg-ef1a-DO-DIOTdTomato-eGFP in the LNAc and labeling neurons in the AIC (fig. s6, C and D). According to these data, despite D1 positive and D1 negative neurons projecting to the LNAc, most neurons are activated by rewarding and aversive social experiences that contain D1Rs.

Next, we examined the Ca^2+^ activity of D1-AIC neurons projecting to the LNAc during rewarding and aversive social experiences using the miniscope (Fig. 3A). Notably, an overall increase in Ca^2+^ activity was observed in D1-AIC neurons projecting to the LNAc in both conditions (Fig. 3, B and C). AIC to NAc projecting neurons showed increased activity during both positive and negative contacts, regardless of proximity (Fig. 3, D to F, fig s7, C and D, fig s8) suggesting that the activity encodes for interaction rather than distance. Moreover, we identified a 67.5% overlap in the neuronal population that was active during both experiences (Fig. 3G). Given the absence of discernible differences in z-score activity between the two experiences, as we previously described, we employed FFT analysis to investigate frequency domain differences (Fig. 3H). Contrary to the LNAc, we observed a shift in activity patterns from lower to higher frequency bands in the AIC when transitioning from rewarding to aversive experiences (Fig. 3I to K). These data suggest that the activity of AIC neurons projecting to LNAc differently encode the rewarding or aversive social experiences.

**Figure 3.**
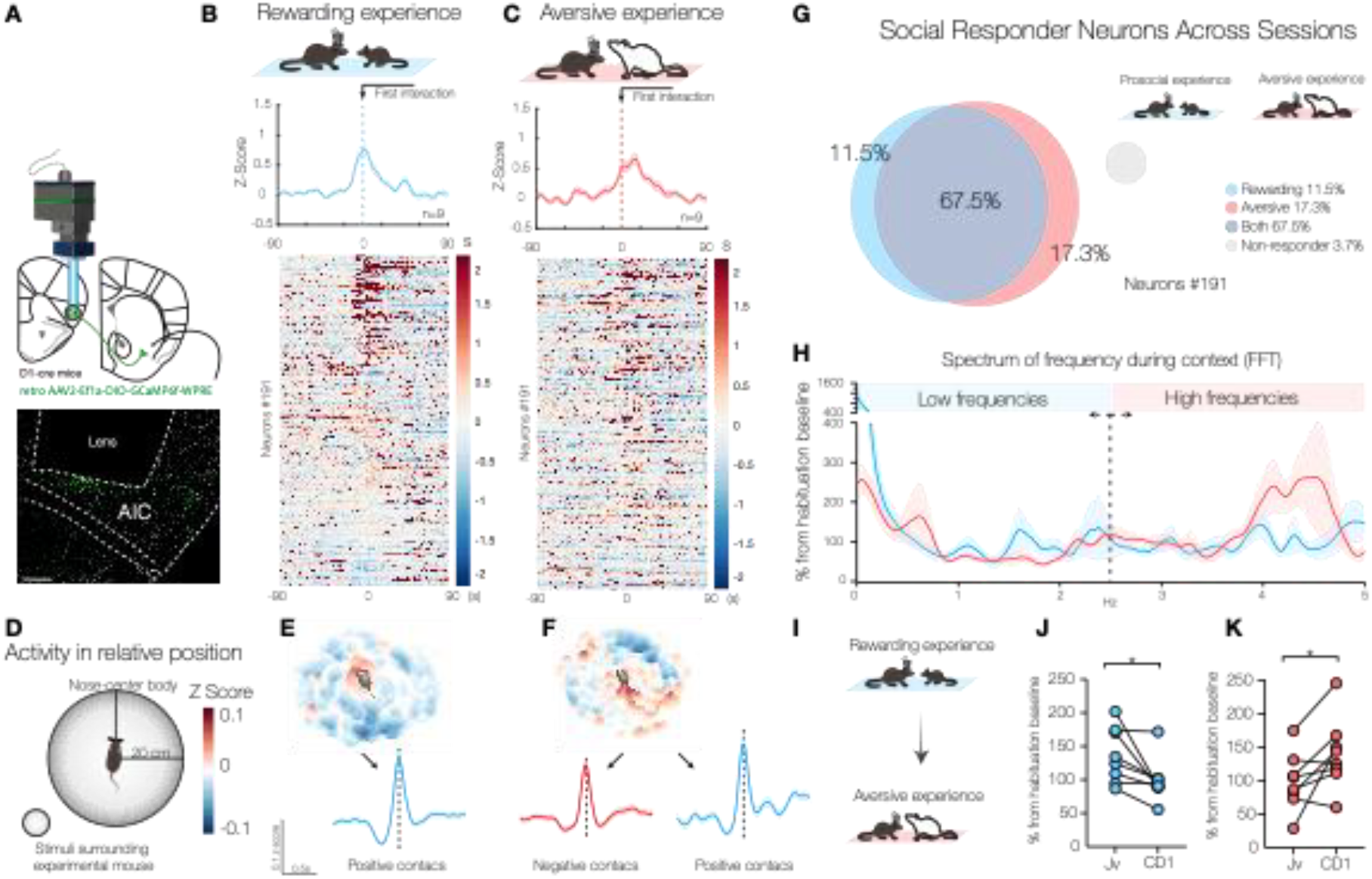
D1R-AIC neurons predicts social valence. (**A**) Schematic of injection and implantation sites. (**B**) Average activity during a rewarding experience aligned with the first interaction. The heatmap represents the activity of individual neurons (each row represents one neuron). (**C**) Average activity during an aversive experience aligned with the first interaction. The heatmap represents the activity of individual neurons (each row represents one neuron). (**D**) Schematic of a 3D heatmap representing the calcium activity in position relative to the stimuli. (**E**) Heatmap showing the activity in positions relative to juvenile stimuli. (**F**) Heatmap showing the activity in positions relative to CD1 stimuli. (**G**) Proportion of neurons that respond to social stimuli. (**H**) Spectrum of frequencies derived from a Fast Fourier Transform (FFT). (**I**) Schematic showing the animals’ first and second exposures. (**J**) Quantification of the low-frequency spectrum between the two exposures (0-2.5Hz). (**K**) Quantification of the high-frequency spectrum between the two exposures (2.5-5Hz). All analysis were performed on the same set of neurons that were tracked across sessions for all panels.

### Valence-dependent synaptic plasticity at AIC to LNAc synapses

Since mice learn to approach or avoid a conspecific depending on their experience (Fig. 1A), and because the AIC to LNAc pathway encodes social valence, we hypothesized that learning occurs through the induction of valence-specific forms of synaptic plasticity at excitatory inputs to the LNAc. Several studies indeed have linked synaptic strengthening with learned behaviors through modulation of AMPA receptors (*12*–*16*). We first verified the specificity of the projection from D1-containing AIC neurons to D1-MSNs in the LNAc. We injected AAV-FLEX-ChR2 in AIC and AAV-FLEX-mCherry in the LNAc in D1-Cre mice (Fig 4A and fig. s9A) and performed double whole cell patch clamp recording from labeled D1 positive and D1 negative neurons in the LNAc (fig. s9B). We found that D1-expressing AIC neurons preferentially projected to D1-MSN in the LNAc, revealing a top-down control (fig. s9, C and D). We then exposed mice to rewarding or aversive social experiences, performed whole-cell recordings from D1-MSNs in the LNAc 24h later while simultaneously recording excitatory inputs from the D1-containing neurons in the AIC (Fig. 4, B and C). We measured the relative ratio of AMPAR to NMDAR mediated currents (AMPA/NMDA ratio) as a proxy of synaptic strength and rectification index of AMPARs to estimate the proportion of calcium-permeable receptors (CP-AMPARs). We observed an increase in the AMPA/NMDA ratio following rewarding social experiences and rectification after aversive social experiences (Fig. 4, D and E). Rewarding or aversive social interactions induce distinct synaptic plasticity at synapses between D1 receptor expressing neurons of the AIC and the LNAc (fig. s10, A to D).

**Fig. 4.**
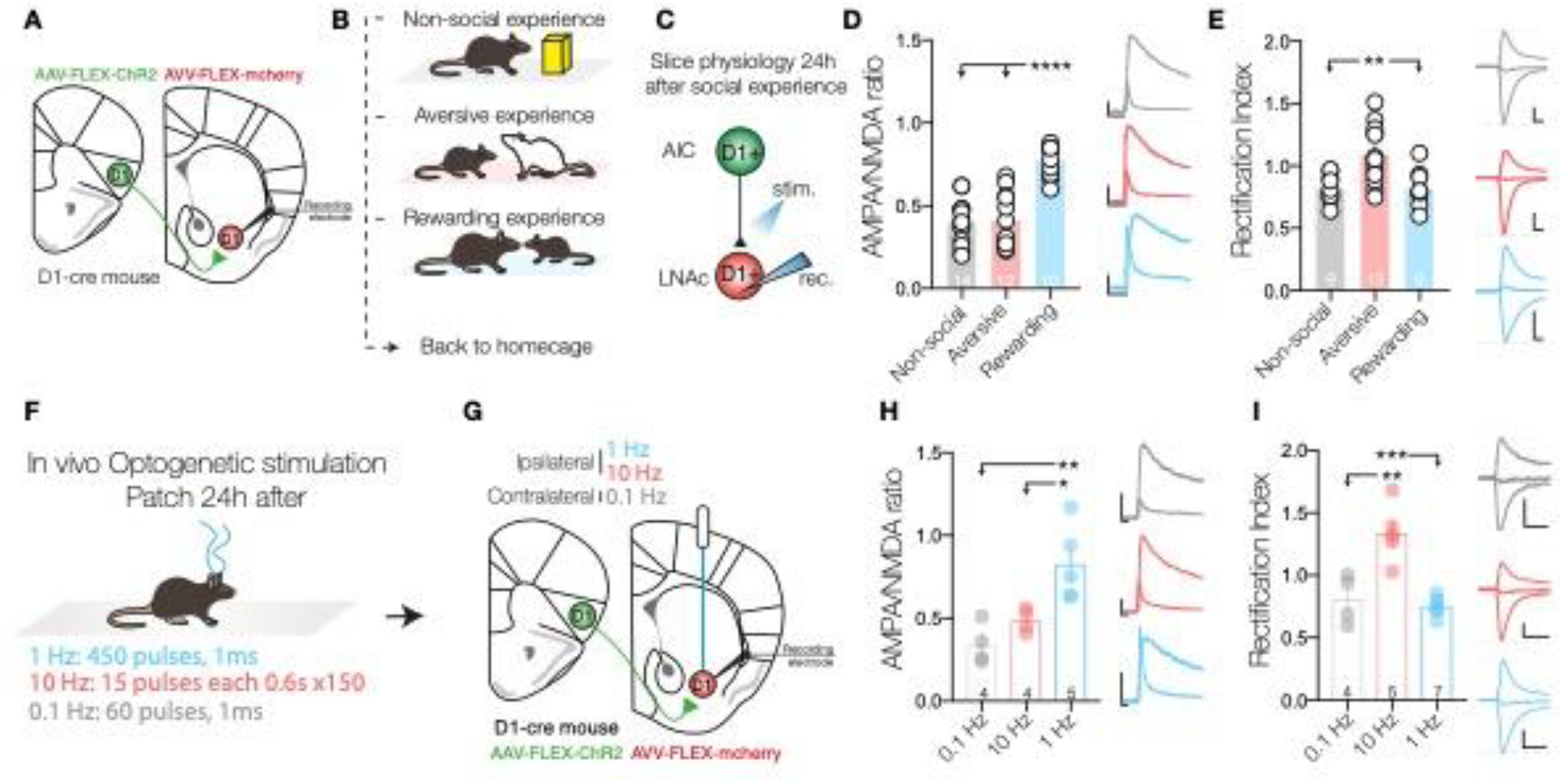
Social valence induce specific long-term synaptic plasticity in AIC-LNAc pathway. (**A**) Schematic representation of viral injection into the Lateral Nucleus Accumbens (LNAc) and Anterior Insular Cortex (AIC). (**B**) Schematic and representative trace of paired recordings between D1-positive and D1-negative Medium Spiny Neurons (MSNs) in the LNAc. (**C**) Diagrammatic representation of the behavioral paradigm. (**D**) Schematic of patch-clam whole cell recording protocol. (**E**) Bar graph showing the AMPA/NMDA ratio 24 hours post-behavioral experiment; to the right are example traces, with **** indicating p < 0.0001. (**F**) Bar graph showing the rectification index 24 hours after free interaction; to the right are example traces, with ** denoting p = 0.0094 for Aversive vs. Prosocial and ** for Aversive vs. Object p = 0.0077. (**G**) Diagram of the in vivo stimulation protocol. (**H**) Schematic representation of viral injection into the Lateral Nucleus Accumbens (LNAc) and Anterior Insular Cortex (AIC). (**I**) Bar graph showing the AMPA/NMDA ratio 24 hours following optogenetic stimulation; to the right are example traces, with ** indicating p = 0.0036 and * p = 0.0292. (**J**) Bar graph of the rectification index; to the right are example traces, with ** p = 0.0011 and *** p = 0.0001. A one-way ANOVA was utilized for statistical testing. Numbers within bars indicate the number of cells in each group. Data are presented as mean ± SEM. Scale bars: 50 pA/10 ms.

Because of the FFT data analysis (Fig. 3, I to K), we then hypothesize that valence-specific activity patterns at the AIC-LNAc pathway induce distinct synaptic plasticity forms during rewarding and aversive interactions. To test this hypothesis, we conducted experiments in D1-Cre mice, injecting AAV-FLEX-ChR2 in the AIC and AAV-FLEX-mCherry in the NAc and bilaterally implanting optic fibers in the LNAc (Fig. 4F). We applied 1 Hz or 10 Hz frequency stimulation to the AIC-LNAc pathway while stimulating the contralateral hemisphere at 0.1 Hz (Fig. 4G). After 24h of optogenetic stimulation, we conducted whole-cell patch clamp recording from D1-MSNs in the LNAc. We observed an increase in AMPA/NMDA ratio after 1Hz stimulation and an increase in rectification index following 10 Hz stimulation (Fig. 4, H and I). As concluded by these data, rewarding and aversive social experiences produce different activity patterns at the AIC and LNAc pathway and as consequence distinct forms of plasticity are elicited.

### Valence-dependent synaptic plasticity controls subsequent approach behavior

Next, we aimed to evaluate the contribution of valence-specific synaptic plasticity at AIC to LNAc synapses in social approach and avoidance learning. We exposed experimental mice to reward social experience and applied 10Hz stimulation at the AIC to LNAc pathway during interaction contacts (Fig. 5, A and B), replicating the pattern observed during aversive social interactions (Fig. 4H). We conducted the approach/avoidance test on the second day (Fig. 5C) and performed whole-cell patch clamp recordings of D1-MSNs in the LNAc while stimulating AIC inputs. In the control group, mice exhibited approach behavior towards the juvenile stimuli during the test phase. However, the approach behavior was abolished entirely in mice exposed to juvenile C57BL/6J and coupled with medium-frequency stimulation during contact (Fig. 5, D and E). Subsequently, we performed electrophysiological recordings from D1-MSNs in the same group of mice. We observed a notable disparity in the AMPA/NMDA ratio between the two groups. Specifically, the ratio recorded after interaction with juvenile C57BL/6J was significantly higher than that observed in mice where contacts were accompanied by 10 Hz stimulation (Fig. 5F). Interestingly, we didn’t find an increase in rectification index (Fig. 5G). However, identified a linear correlation between the rectification index and the avoidance index: a higher proportion of GluA2-lacking AMPARs at the AIC-LNAc synapses corresponded to a greater degree of avoidance behavior towards the juvenile stimulus (Fig. 5H).

**Fig. 5.**
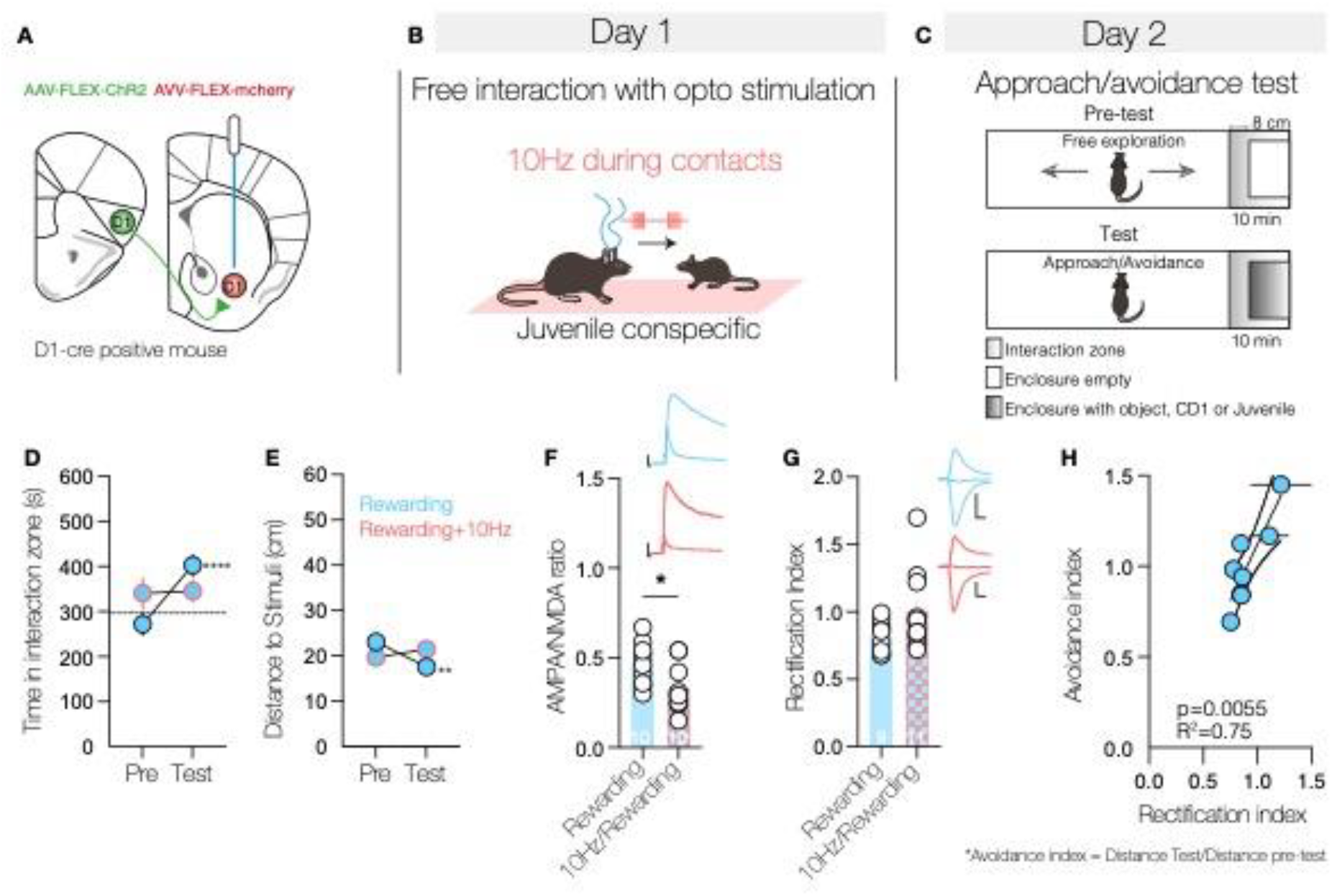
Valence-dependent synaptic plasticity is essential for learned approach behavior. (**A**) Schematic indicating the site of viral injection and placement of the optical fiber. (**B**) Diagram detailing the protocol, and the delivery of 10Hz optical stimulation during bouts of contact. (**C**) On Day 2, animals were placed in a new arena and, after 10 minutes of free exploration with an empty enclosure, were introduced to an enclosure containing the stimuli from the previous day. (**D**) Graph depicting the duration of interaction within the zone, with **** signifying p < 0.0001 for animals that did not receive the stimulation and p = 0.8745 for animals that received the 10Hz stimulation. (**E**) Graph depicting the distance to stimuli, with ** indicating p = 0.0027 and p = 0.4030 for rewarding experiences with 10Hz stimulation. (**F**) Bar graph showing the AMPA/NMDA ratio, with * indicating p = 0.020. (**G**) Bar graph showing the rectification index, p = 0.1048. (**H**) A correlation graph between the rectification index and the avoidance index, calculated as the distance during the post-test divided by the distance during the pre-test, with p = 0.0055 and R^2^ = 0.75 (each dot represents a single animal, n = 8). A paired t-test was utilized for analyses in (D) and (E). An unpaired t-test was employed for (F) and (G). The numbers inside the bars denote the number of cells in each group. Data are presented as mean ± SEM. Scale bars: 50 pA/10 ms.

In this study, we characterize a novel top-down circuit between D1 neurons in the AIC and D1 neurons in LNAc, which undergoes forms of social valence-dependent synaptic plasticity. These forms of plasticity serves as the neuronal substrate for learned approach and avoidance behaviors. Interestingly, these forms of plasticity arise from distinct frequency patterns, defining a specific signature for social valence. It is important to note that in this context, valence encompasses more than just the activation or inhibition of a specific neuronal subpopulation and deviates from the traditional definition (*17, 18*). Our data indicate that valence is determined by the frequency patterns within a specific pathway and the subsequent experience-dependent synaptic changes that drive learned behaviors.

## Acknowledgments

We thank Lorena Jourdain for her assistance with the experimental procedures and Giuseppe Chindemi for his help with data analysis, which unfortunately we could not include in the final version of this publication. We also thank Christian Lüscher for his suggestions on the manuscript.

## Funding

P.E. is supported by the Swiss Government Excellence Scholarship (FCS) for PhD Studies Grant ESKAS-Nr: 2017.0922. C.B. is supported by the Swiss National Science Foundation and by the ERC SocialNac.

## Author Contributions

P.E. and C.B. conceived the project. P.E. and C.B. wrote the manuscript. P.E. conducted all the experiments, with F.C. assisting with part of the patch-clamp experiments. B.G. performed the behavioral analyses using DeepLabCut (DLC) and SimBA, as well as the analysis of calcium activity relative to contacts and position. P.E. carried out the anatomical tracing experiments with the assistance of M.L. Manual cell counting was performed by M.L. and V.T. P.E. conducted the analyses and statistical evaluations. Figures were prepared by P.E.

## Competing interests

The authors declare no competing interests.

## Data and materials availability

Scripts and raw data will be uploaded to a server to grant access to them.

## Notes

### Competing Interest Statement

The authors have declared no competing interest.

### Summary of Updates

We have revised the text and the figures for submission to another journal

